# Rapid three dimensional two photon neural population scanning

**DOI:** 10.1101/029090

**Authors:** Renaud Schuck, Peter Quicke, Caroline Copeland, Stefania Garasto, Luca A. Annecchino, June Kyu Hwang, Simon R. Schultz, EMBS Member

## Abstract

Recording the activity of neural populations at high sampling rates is a fundamental requirement for understanding computation in neural circuits. Two photon microscopy provides one promising approach towards this. However, neural circuits are three dimensional, and functional imaging in two dimensions fails to capture the 3D nature of neural dynamics. Electrically tunable lenses (ETLs) provide a simple and cheap method to extend laser scanning microscopy into the relatively unexploited third dimension. We have therefore incorporated them into our Adaptive Spiral Scanning (SSA) algorithm, which calculates kinematically efficient scanning strategies using radially modulated spiral paths. We characterised the response of the ETL, incorporated its dynamics using MATLAB models of the SSA algorithm and tested the models on populations of Izhikevich neurons of varying size and density. From this, we show that our algorithms can theoretically at least achieve sampling rates of 36.2Hz compared to 21.6Hz previously reported for 3D scanning techniques.

## I. INTRODUCTION

Understanding neuronal dynamics is a key step towards comprehending brain function and dysfunction and will help inform future treatments of brain disorders. In order to understand these dynamics the activity of many neurons must be sampled at high frequency and in three dimensions. Multiphoton laser scanning microscopy (MPLSM) of genetically encoded Ca^2+^ indicators enables high spatial (*µ*m) and rapidly improving temporal (ms) resolution imaging of large numbers (up to 1000) of labelled neuronal subtypes. Thus far most imaging occurs in two dimensions due to the difficulty of quickly shifting the focal plane along the optical axis. This precludes cross-layer cortical information flow capture. To address this problem, several strategies evolved to image in 3D including acousto-optic devices (AODs) [1], mechanical objective translation [2], wavefront shaping [3] and electrically tunable lenses (ETLs) [4]. All of these techniques except ETLs require custom built or extensive additions to microscopes, whereas an ETL can easily integrate into a commercial MPLSM setup. Previous implementations combined ETLs with x-y raster scanning; however, as demonstrated by our previous work [5], ‘smart’ scanning strategies increase sample rates 5-10x by reducing time spent scanning extracellular space. Our algorithms take galvanometric scanners (GSs) and ETL inertia into account to calculate a faster spiral scanning trajectory in 3D. In this paper, we show two examples of 3D scanning strategies that resolve Ca^2+^ transients from GCaMP6f for a population of 800 neurons.

## II. MATERIALS AND METHODS

### A. ETL Model

Our system uses an ETL (EL-10-30-VIS-LD, Optotune) in conjunction with an offset lens (f = -75mm, LC4513-A, Thorlabs) placed close to the objective rear aperture, powered by a current source (LD3000R, Thorlabs), to axially translate the focal plane. We measured the current source command voltage/axial shift curve (Figure 1) using a phantom of 1*µ*m fluorescent beads (F-13080, Life Technologies) suspended in 1% agarose. By measuring the objective displacement required to refocus the microscope onto beads as a function of voltage, we show that the ETL can be approximated by a linear system over a range of 120-800 *µm*. This range spans up to layer five of the mouse cortical column. Figure 1 shows the data with a linear fit with a slope of -3508*µ*mV^−1^ and an offset of 51.94*µ*m.

**Fig. 1:**
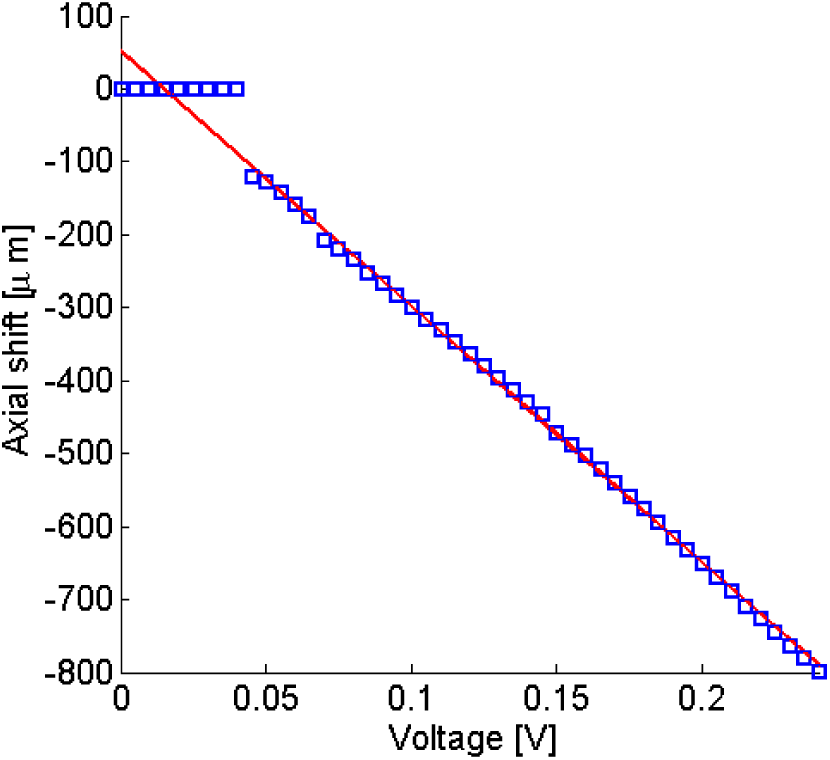
The measured axial shift of the focal plane (blue squares) as a function of command voltage fit with a linear function (red line) which we use to calculate its response. A = *f* (V) = −3508V + 51.94 where A is the axial shift in microns and V is the command voltage.

Figure 2 shows ETL step response data measured by Optotune [6] fit with our model (see table I), a damped sine modulated by a first order exponential:

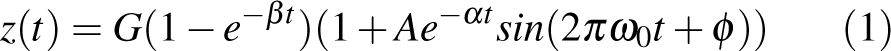

**Fig. 2:**
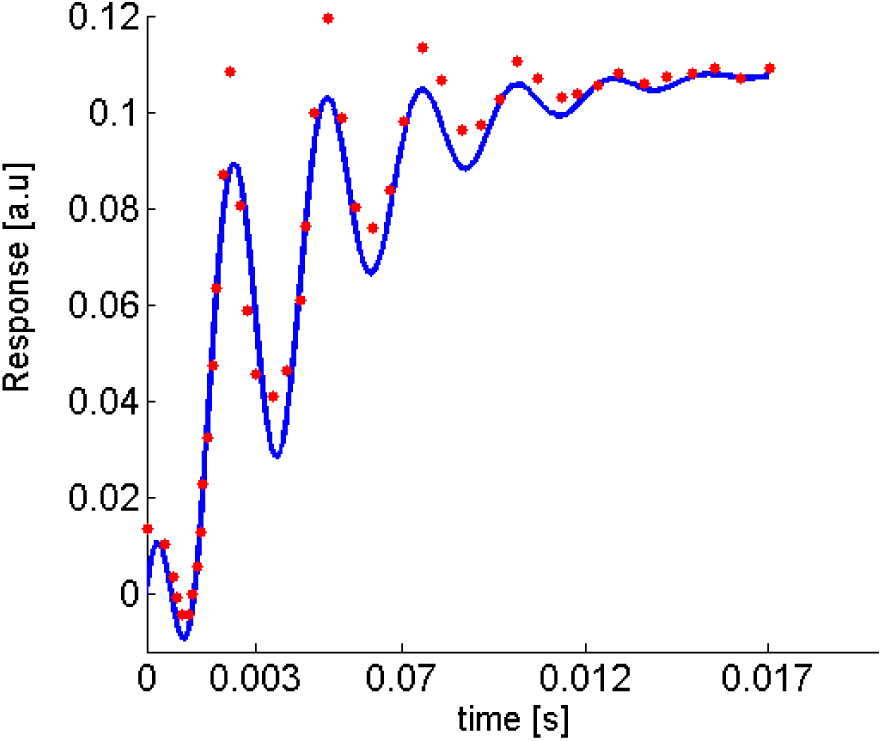
Our model of the dynamic response of the ETL (blue line) fit to the step response recorded by Optotune (red points).

**TABLE I:** Fitted parameters for our model of the ETL step response function. G is the gain, *β* the time constant of the first order exponential, *ω*_0_ the natural oscillation frequency, *A* the amplitude of the oscillation, *α* the damping factor and *φ* the phase.

| $G$ [mm] | $\beta$ [$s^{-1}$] | $A$ [mm] | $\alpha$ [$s^{-1}$] | $\omega_0$ [Hz] | $\phi$ [rad] |
| --- | --- | --- | --- | --- | --- |
| 0.1096 | 243 | -2.011 | 357.2 | 384.4 | -0.9332 |

### B. Scanning and neural population model

Our previous work introducing the 2D SSA algorithm [5] showed how we characterised the galvanometer’s (6215H, Cambridge Technology) transfer function similarly to the ETL. In order to validate the scanning algorithm on real brain tissue samples, GFP was expressed in mouse primary somatosensory cortex (S1) via stereotactic injection of adenoassociated viruses under the CAG promoter into C57BL/6 mice (Figure 3). 6 weeks post-injection, animals were perfused with 4% PFA, the brain removed and submerged in ACSF prior to 2-photon imaging. Procedures were Home Office (UK) approved and accorded with the Animals (Scientific Procedures) Act 1986. 3D algorithms use the transfer functions to calculate the driving signals, *I_X_*, *I_Y_* and *V_z_* delivered to the two GSs and the ETL respectively, to move the focal point to the coordinates *X*,*Y*,*Z*. The algorithm, given a set of neuronal coordinates, sorts the neurons by distance from the center of the field of view (FOV) and increasing axial depth. Starting at the FOV center it scans the neurons in an x-y spiral trajectory in a z-plane defined by the center of the first neuron. Once the spiral reaches the first neuron’s center the axial focal depth is increased stepwise using the ETL. Neuron densities are usually high enough such that the spiral will have sampled activity from the tops of deeper neurons. Therefore the depth is increased to coincide with the shallowest neuron center not already sampled sufficiently by the beam. This is shown schematically in figure 4. To test the algorithm we generated surrogate data from functional fluorescent signals governed by calcium indicator GCaMP6f dynamics [7] from firing patterns of Izhikevich neurons [8], [5].

**Fig. 3:**
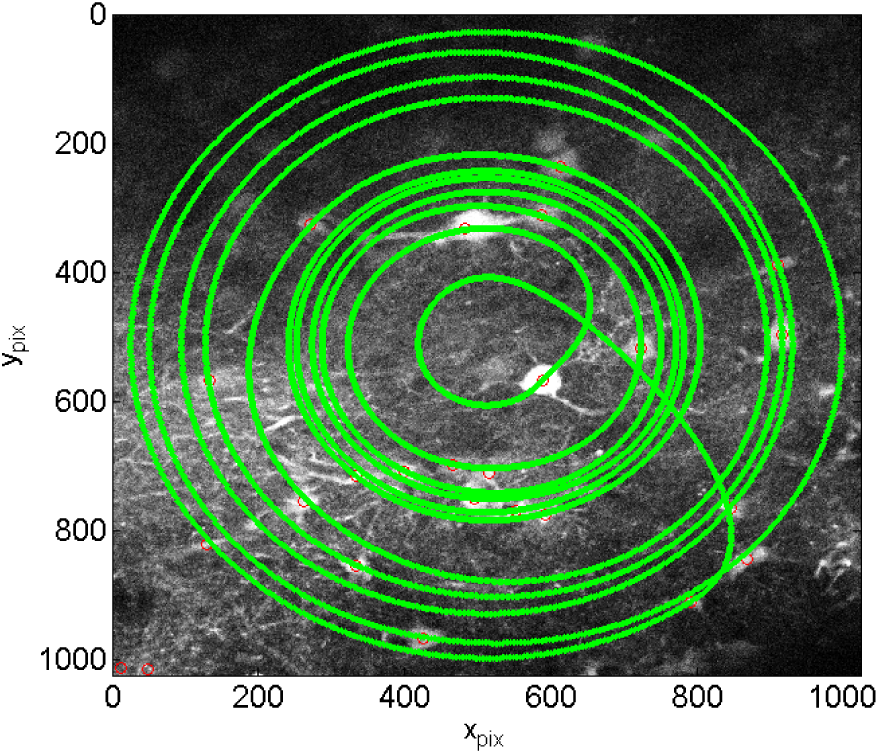
An example beam path from our SSA algorithm applied to GFP labelled neurons. We acquired a full frame raster scan of 1024x1024 pixels and automatically extracted the coordinates of the neuron centres (red circles). Finally, we used these coordinates with our 2DSSA algorithm to generate a spiral scanning path (green line).

**Fig. 4:**
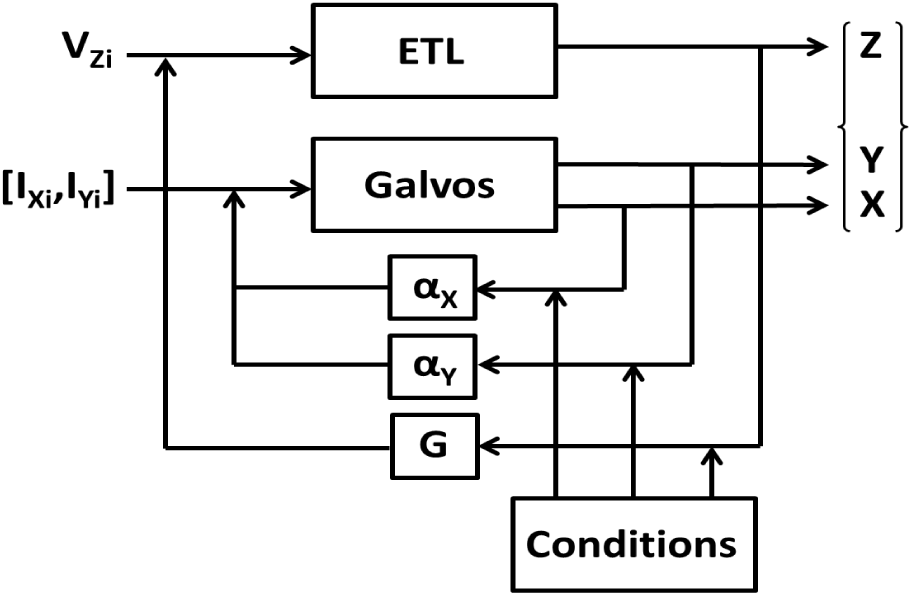
3D Scanning model block diagram. *α_X_*, *α_Y_* and *G* are the gains in the open loop from *GS_X_*, *GS_Y_* and the ETL respectively which are modulated to produce axially displaced spiral paths which sufficiently sample all neurons.

## III. RESULTS

Combining a simulated population of neurons emitting fluorescent signals and a 3D dynamical model of the scanning elements from a MPSLM, we built a platform that is able to test various kinds of scanning strategies. Here we describe the performance and limitations of the SSA algorithm extension.

### A. SSA Algorithm extension for 3D scanning

We used the SSA algorithm in a 2D experiment on 6*µm* fluorescent beads (Polyscience Inc) suspended in 1% Agarose. Driving the GSs above a threshold frequency elicited position drift. Hence, in our simulation we decreased the amplitude modulated sine frequency applied to the GSs *I_X_*, *I_Y_* from *ω_I_* = 10*kHz* to *ω_I_* = 1*kHz*. Two 3D Algorithm were tested: the ‘Cylindrical’ scanning strategy (CST) and

**Algorithm:**
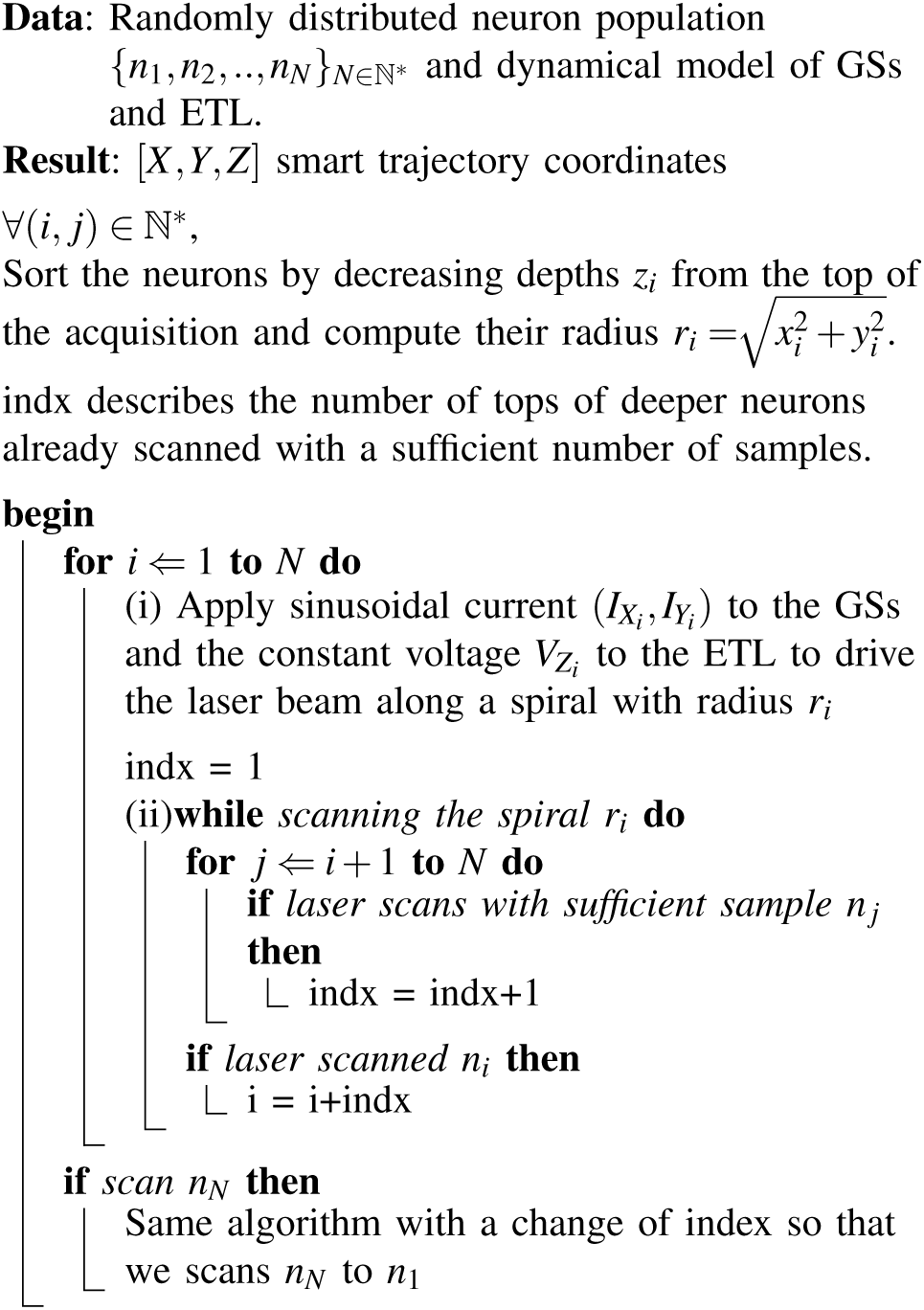
3D Cylinder Scanning Strategy.

‘Orbital’ scanning strategy (OST) which differ by the type of ETL driving signal. For the CST, the voltage *V_z_* applied to the ETL is a series of steps: 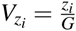 where *i* ∈{1..*N*}, *G* is the gain of the ETL, [*z_i_*]*_i_*_∈{1.._*_N_*_}_ are the planes occupied by the neurons and N is the total number of scanned cells (See Algorithm (i)). For the OST, the voltage *V_z_* applied to the ETL is an amplitude modulated sinusoid: 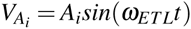 where *A_i_*_∈{1.._*_N_*_}_ is the amplitude and *ω_ETL_* the frequency of the voltage applied to the ETL. For reasons of clarity, we present only the CST algorithm in detail here. We combined the previously CST presented driving signals with specific geometric conditions (see Algorithm (ii) and Figure 4) resulting in modulated, tilted, circular trajectories fit to a randomly positioned 3D set of cells (Figure 5*A*, 5*B* and 5*C*), reducing the impact of the GSs inertia and where the user can define a neuron dwell time determining the time spent in each axial plane.

**Fig. 5:**
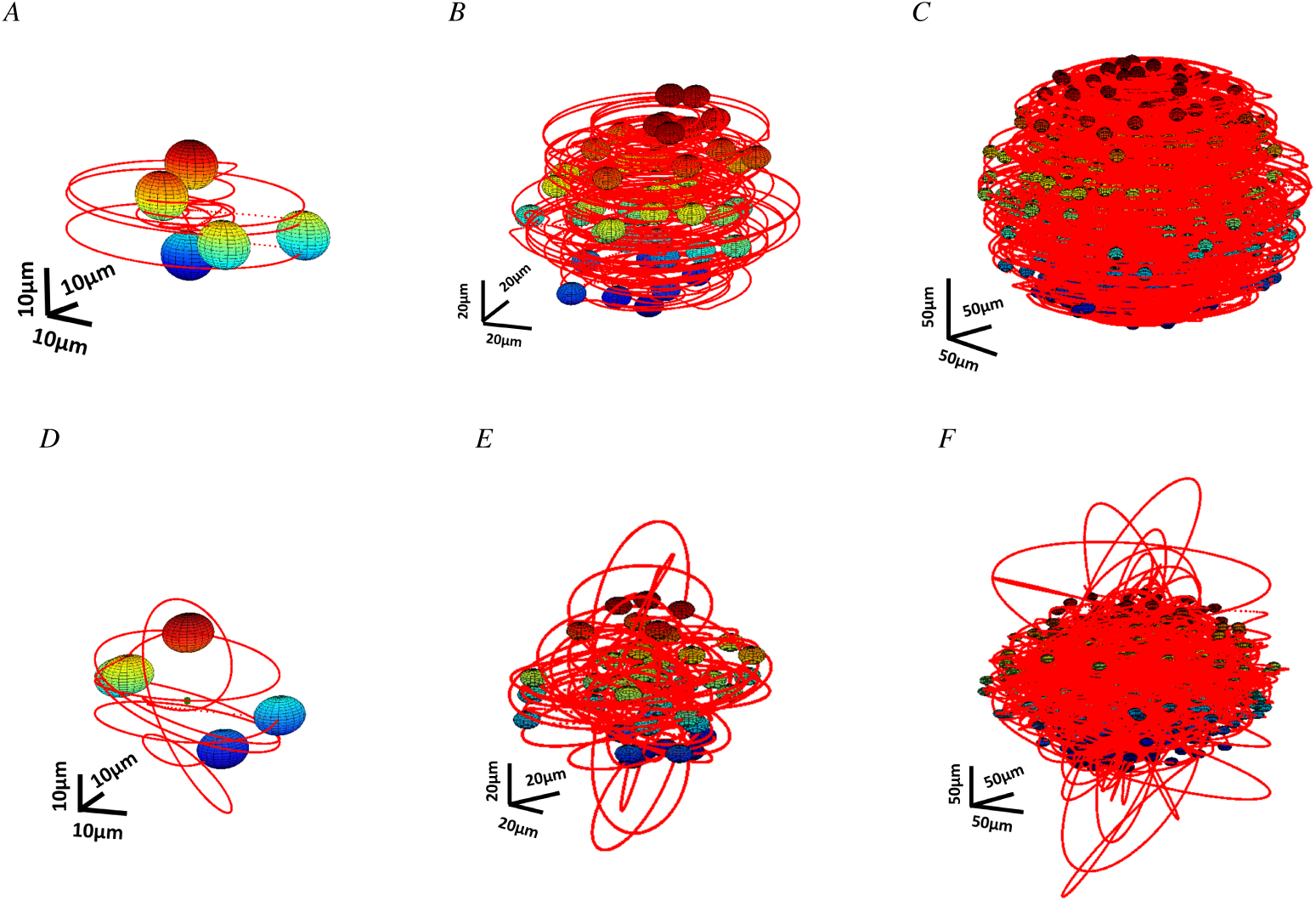
Two examples of 3D MPLSM scanning strategies on pseudo-random population of neurons. A, B and C represent the laser path in red for the CST for five, fifty and five hundred cells. The spherical neuron distribution is modelled with the spatial density of active cells in mouse V1 [9]. D, E and F represent the laser path of the OST.

### B. Sampling frequency and Calcium transient acquisition

Driving the dynamics of the GSs and the ETL with the above described driving signals enabled us to calculate the sampling frequency of our 3D scanning strategies across various neural geometries. Figure 6*A* shows that for a constant density of cells, there is a log-log linear decrease in sampling frequency for both the CST and the OST. Theoretically, single neuron action potential extraction necessitates a sampling rate over the Nyquist frequency corresponding to a calcium transient from a specific fluorescent protein. Based on these requirements, our simulation simultaneously resolved calcium transients (GCaMP6f) from 800 neurons with the CST and 700 neurons with the OST (Figure 6*A*). Our simulated raw PMT signal acquisition is based on the same principle as for the 2DSSA [5]. Figure 6*B*, 6*C* and 6*D* show ground truth fluorescent (green) and reconstructed Calcium transient (blue) from 5, 50 and 100 neurons sampled at 156*Hz*, 18*Hz* and 10*Hz* respectively for the CST.

**Fig. 6:**
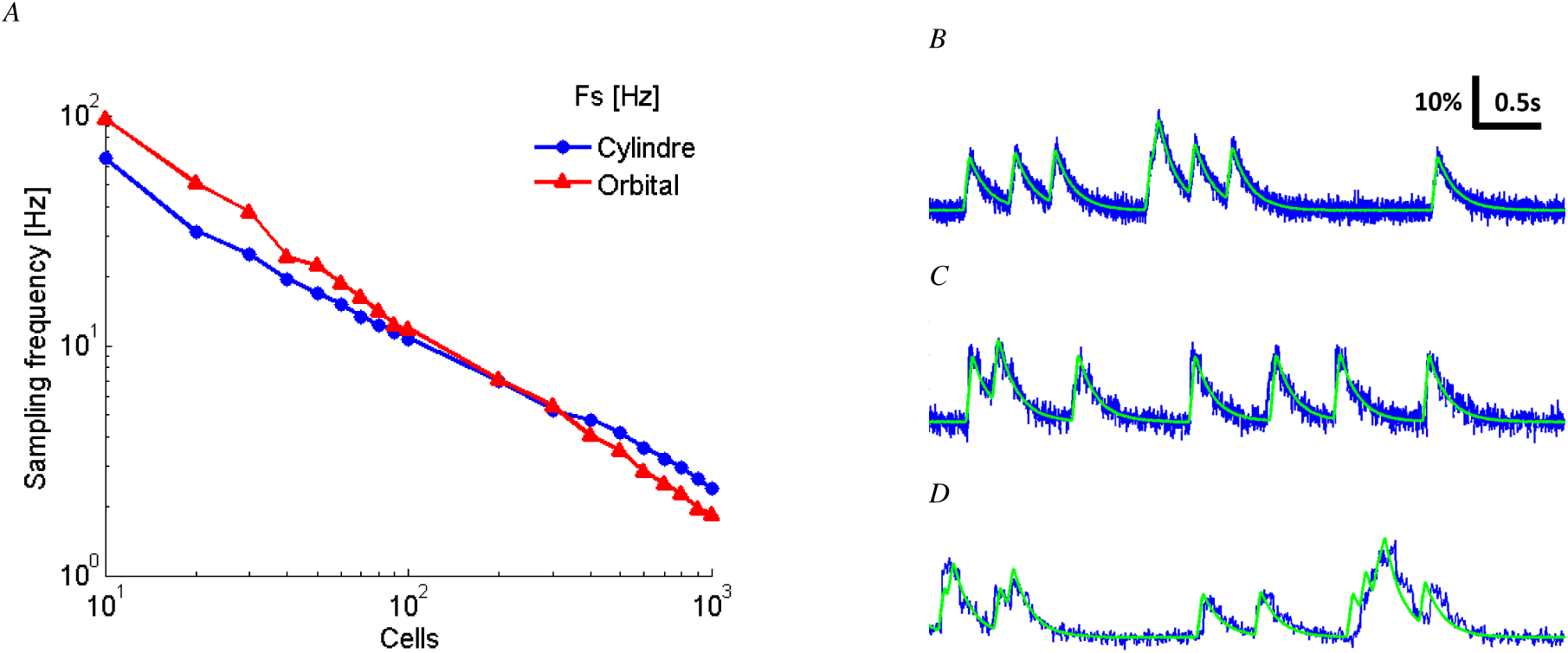
3DSSA allows the sampling of high number of neurons. A is a loglog plot representing the Sampling frequency decay of both CST and OST according to the number of cells. This was done for a specific neuron density *d* = 9.5 ×10^−5^*µm*^−3^ and shows that we can resolve calcium transient for up to 800 cells with the cylinder scanning strategy. B, C and D describe ground truth fluorescent in green and reconstructed Calcium transient in blue from 5, 50 and 100 neurons sampled at 156*Hz*, 18*Hz* and 10*Hz* over 5*s*, respectively with the CST.

## IV. DISCUSSION

Here we present faster 3D scanning strategies using an ETL. Previous studies such as [4] reported a 21.6Hz scan rate for 34 neurons in two planes separated by 40*µ*m using AODs. Our CST can sample 40 neurons distributed in and between equivalent planes using cheaper GSs at a higher rate of 36.2Hz. Simulations also indicate our strategy will be able to sample 800 neurons sufficiently fast to detect Ca^2+^ transients from GCaMP6f. Our OST further increases scanning speeds up to 51.6Hz for the same situation and offers significant increases for up to 200 neurons (Figure 6). We also hope to further increase scan speeds by applying optimal control theory to our 3D algorithms to find kinematically optimal modulations of the driving signals applied to GSs and ETL. The highest scanning speed that we can achieve whilst maintaining high enough sampling for good spike detection accuracy [10], [11] remains to be seen. One possible problem we anticipate when we implement high speed scanning in hardware is drift in focal spot position caused by heating of the ETL and GSs. This could cause the trajectory to drift enough to bypass some neurons completely and require software compensation. We are currently working towards validating our model by imaging functional fluorescent indicators in mouse cortex both *in vivo* and *in vitro*.

## V. ACKNOWLEDGEMENTS

We thank Stephan Smolka of Optotune for useful discussions on ETL technology.

## Notes

This work was supported by EU FP7 Marie Curie Initial Training Network 289146, a Royal Society Industry Fellowship to SRS, BBSRC grant BB/K001817/1 to SRS, and by Scientifica Ltd.

## REFERENCES

[1] G. Katona, G. Szalay, P. Maák, A. Kaszás, M. Veress, D. Hillier, B. Chiovini, E. S. Vizi, B. Roska, and B. Rózsa, “Fast two-photon in vivo imaging with three-dimensional random-access scanning in large tissue volumes,” Nature methods, vol. 9, no. 2, pp. 201–208, 2012.

[2] W. Göbel, B. M. Kampa, and F. Helmchen, “Imaging cellular network dynamics in three dimensions using fast 3d laser scanning” Nature methods, vol. 4, no. 1, pp. 73–79, 2007.

[3] S. Quirin, J. Jackson, D. S. Peterka, and R. Yuste, “Simultaneous imaging of neural activity in three dimensions” Frontiers in neural circuits, vol. 8, 2014.

[4] B. F. Grewe, F. F. Voigt, M. vant Hoff, and F. Helmchen, “Fast two-layer two-photon imaging of neuronal cell populations using an electrically tunable lens” Biomedical optics express, vol. 2, no. 7, pp. 2035–2046, 2011.

[5] R. Schuck, L. A. Annecchino, and S. R. Schultz, “Scaling up multiphoton neural scanning: The SSA algorithm” in Engineering in Medicine and Biology Society (EMBC), 2014 36th Annual International Conference of the IEEE. IEEE, 2014, pp. 2837–2840.

[6] Datasheet: EL-10-30-Series, Optotune AG, 2014, updated: 13.04.2015.

[7] T.-W. Chen, T. J. Wardill, Y. Sun, S. R. Pulver, S. L. Renninger, A. Baohan, E. R. Schreiter, R. A. Kerr, M. B. Orger, V. Jayaraman et al., “Ultrasensitive fluorescent proteins for imaging neuronal activity” Nature, vol. 499, no. 7458, pp. 295–300, 2013.

[8] E. M. Izhikevich et al., “Simple model of spiking neurons” IEEE Transactions on neural networks, vol. 14, no. 6, pp. 1569–1572, 2003.

[9] R. Cossart, D. Aronov, and R. Yuste, “Attractor dynamics of network up states in the neocortex” Nature, vol. 423, no. 6937, pp. 283–288, 2003.

[10] J. Onativia, S. R. Schultz, and P. L. Dragotti, “A finite rate of innovation algorithm for fast and accurate spike detection from two-photon calcium imaging” Journal of neural engineering, vol. 10, no. 4, p. 046017, 2013.

[11] S. Reynolds, J. Oñativia, C. S. Copeland, S. R. Schultz, and P. L. Dragotti, “Spike detection using FRI methods and protein calcium sensors: performance analysis and comparisons” in 11th international conference on Sampling Theory and Applications (SampTA 2015), Washington, DC, USA, May 2015.

